# Highly sensitive and scalable time-resolved RNA sequencing in single cells with scNT-seq2

**DOI:** 10.1101/2025.06.03.657745

**Authors:** Qi Qiu, Hongjie Zhang, William Gao, Fan Li, Dongming Liang, Hao Wu

## Abstract

Understanding gene expression dynamics requires resolving newly synthesized RNAs from pre-existing pools at single-cell resolution. Here, we present scNT-seq2, a highly sensitive and scalable method for time-resolved single-cell RNA sequencing. By systematically optimizing the second-strand cDNA synthesis (2^nd^ SS) step, we substantially improved read alignment rates, reduced background mutations, and enhanced library complexity compared to the original scNT-seq ^1^. Benchmarking in 4sU-labeled K562 cells demonstrated that scNT-seq2 accurately quantifies newly synthesized transcripts and preserves the gene level RNA turnover. The enhanced sensitivity enables robust detection of dynamic, cell-cycle state specific genes, such as S-phase regulators. Together, scNT-seq2 provides an efficient and versatile tool for dissecting transcriptional dynamics across diverse biological systems at single-cell resolution.

## Main

Metabolic RNA labeling offers a powerful strategy for dissecting RNA dynamics by distinguishing newly synthesized (“new”) from pre-existing (“old”) transcripts. This is achieved by incorporating nucleoside analogs such as 4-thiouridine (4sU) or 5-ethynyluridine (5-EU) into nascent RNAs, followed by biochemical enrichment or enrichment-free chemical conversion (e.g. T-to-C substitutions for 4sU labeling) ^2-4^ to selectively identify labeled transcripts. When combined with single-cell RNA-seq platforms – including plate-based [scSLAM-seq^5^; NASC-seq^6,7^; scEU-seq^8^], combinatorial indexing [sci-fate ^9,10^] or droplet-based [scNT-seq ^1^] – this strategy enables the simultaneous profiling of new and old RNAs within individual cells. By offering precise temporal control over metabolic RNA labeling, these methods improve RNA velocity analyses to predict cell state transitions^1,10,11^, enable quantitative inference of RNA kinetics in a specific cell type/state, and enhance gene regulatory network (GRN) analysis ^1,8^, thereby providing a powerful framework for analyzing cell-type-specific transcriptional dynamics^12,13^.

Despite these advances, current single-cell time-resolved RNA-seq methods face a trade-off between sensitivity (i.e., the number of genes detected per cell) and scalability (i.e., number of cells assayed). Plate-based methods including scSLAM-seq and NASC-seq offer higher sensitivity and library complexity but are constrained by low throughput and high cost. In contrast, combinatorial indexing-based approaches such as sci-fate are more scalable but often suffers from reduced sensitivity. Meanwhile, droplet-based methods such as scNT-seq offers a balanced performance of both detection sensitivity and scalability by leveraging high-throughput droplet microfluidics ^14^ and efficient on-bead Timelapse chemical conversion ^3^.

To address this limitation, incorporation of a randomly primed second-strand cDNA synthesis reaction (2^nd^ SS) step after reverse transcription has been shown to enhance library complexity in both single-cell RNA-seq^15^ and spatial transcriptomic platforms (e.g., Slide-seqV2^16^). In our previous work^1^, we demonstrated that introducing a Klenow DNA polymerase/N9 random primer-based 2^nd^ SS reaction into scNT-seq, following chemical conversion and reverse transcription, significantly increased library complexity, likely due to recovery of truncated cDNAs caused by chemical conversions. However, despite its broad utility, 2^nd^ SS reaction conditions have not been systematically optimized across applications.

In this study, we present scNT-seq2, an improved time-resolved single-cell RNA-seq method that achieves enhanced sensitivity by systematically optimizing the second-strand cDNA synthesis step. Key improvements include replacing Klenow polymerase with Bst3 and redesigning the random primer to a N3G2N4B configuration (**Fig. 1a** and **Supplementary Tables 1-2**). Compared to the original scNT-seq protocol, scNT-seq2 exhibits substantially improved read alignment rates, reduced background mutations, and improved 4sU conversion efficiency, along with robust gains in library complexity. Together, these enhancements establish scNT-seq2 as a highly sensitive and scalable platform for time-resolved transcriptomic profiling in single cells.

**Figure 1.**
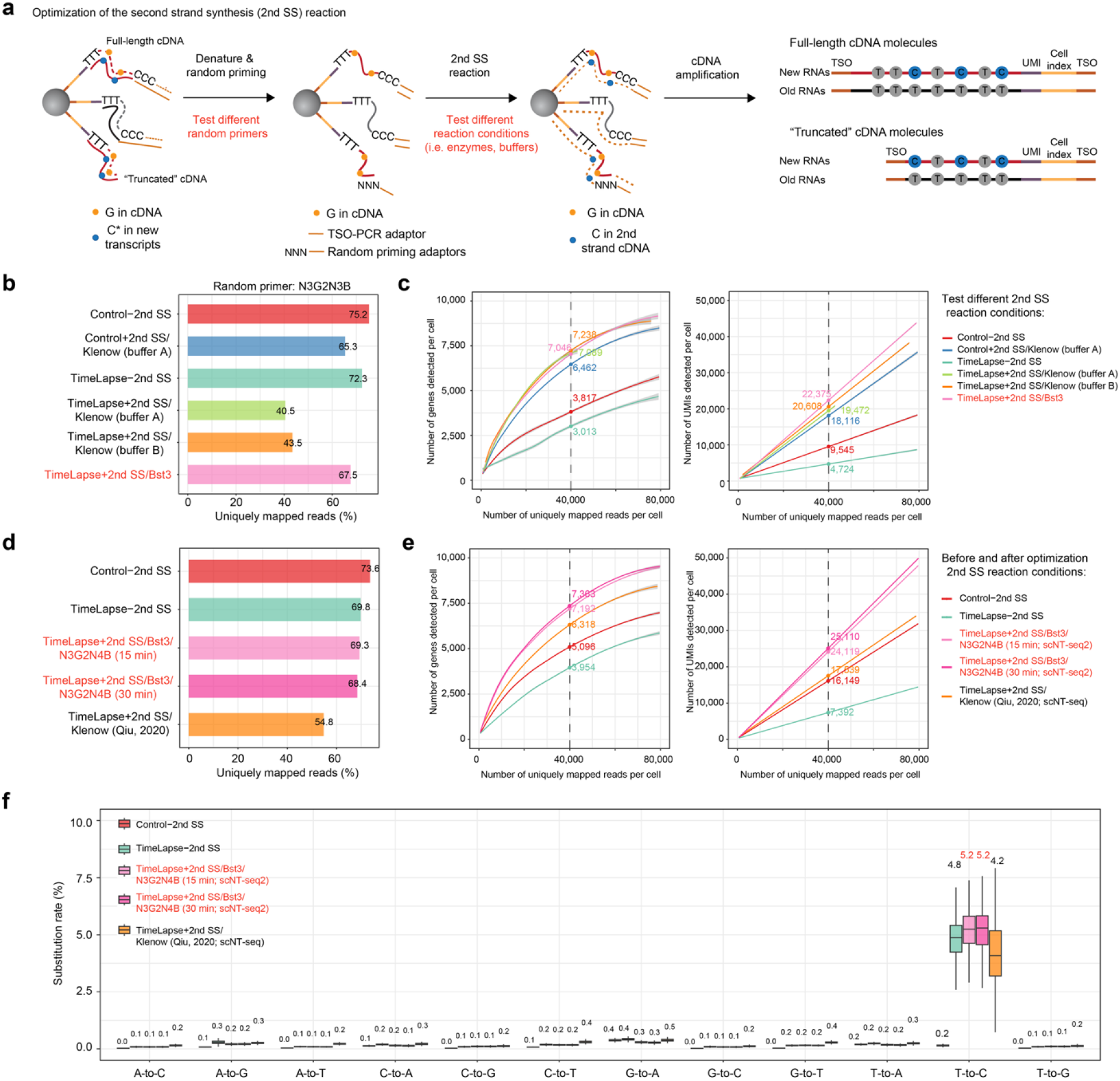
Optimizing second-strand cDNA synthesis reaction in K562 cells. **a**. Schematic representing the experimental strategy for optimizing the 2^nd^ SS reaction in scNT-seq. **b**. Bar graph showing the fraction of uniquely mapped reads various enzymatic reaction conditions. **c**. Fitted line plots illustrating library complexity by comparing genes (left) or UMIs/transcripts (right) detected per cell as a function of aligned reads per cell across various enzymatic reaction conditions. Either reaction buffer A ^15^ or B ^1^ was used with Klenow Fragment (3’->5’ exo-); *Bst* 3.0 DNA polymerase (denoted as Bst3) was identified as the more optimal condition in this study and was used in scNT-seq2. All experiments were performed using the same batch of *in vitro* 4sU-labeled K562 cells (100 μM, 4 hours). Estimated numbers of genes or UMIs detected per cell at matching sequencing depth (40,000 reads per cell) for different experiments are shown. The shaded regions depict 95% confidence intervals. **d**. Bar graph showing the fraction of uniquely mapped reads from the experiments either using optimized 2^nd^ SS or the original Klenow-based 2^nd^ SS reaction. **e**. Fitted line plots illustrating library complexity by comparing genes (left) or UMIs/transcripts (right) detected per cell as a function of aligned reads per cell across experiments in **(d)**. **f**. Box plot comparing nucleotide substitution rates in 4sU-labeled K562 cells from experiments in **(d)**.

### Optimizing second-strand cDNA synthesis using scNT-seq platform

Although incorporating a Klenow/random primer (N9)-based 2^nd^ SS step substantially increase the library complexity of the scNT-seq library^1^, we observed a notable reduction in read alignment rates, from 72.5% (TimeLapse-2^nd^ SS) to 54.8% (TimeLapse+2^nd^ SS/Klenow, **Supplementary Fig. 1a-b**) in K562 cells. Interestingly, 2^nd^ SS alone caused only a minor change in alignment rate, suggesting that the combined effects of on-bead chemical conversion (TFEA/NaIO4 treatment) and 2^nd^ SS contribute to the overall decrease. Furthermore, we detected elevated background mutation rates with the inclusion of 2^nd^ SS.

To address these limitations, we hypothesized that both the design of the random primer and the choice of polymerase/reaction conditions would be critical for optimizing 2^nd^ SS performance (**Fig. 1a**). First, we designed a panel of random primers with varying numbers of guanines (G), random mixed or degenerate bases (N: 25%A, 25%C, 25%G, 25%T; B: 0%A, 33%C, 34%G, 33%T, **Supplementary Table 2**). The inclusion of guanines was intended to enhance primer affinity to the “CCC” sequence at the 5′ end of full-length cDNAs, a sequence derived from the template-switching oligo used during reverse transcription. Second, we tested three reaction conditions including two using Klenow polymerase and one using *Bst* 3.0 DNA polymerase (hereafter, Bst3).

We first evaluated how random primer sequence affects Klenow-based 2^nd^ SS performance in 4sU-labeled K562 cells. Consistent with our previous results, the 2^nd^ SS reaction increased the library complexity but decreased read alignment rate (**Supplementary Fig. 1c-d**). While the library complexity was similar across all primers tested, alignment rates varied considerably, from 40.5% with the N3G2N3B primer to 54.6% with the N3G2N4B primer (**Supplementary Fig. 1c**). Based on this, we selected the N3G2N4B primer for subsequent optimization.

Next, we compared different DNA polymerase reaction conditions with the same random primer (N3G2N3B) in K562 cells. Notably, the Bst3-based condition yielded a substantially higher alignment rates (67.5%) compared to the two Klenow-based conditions (40.5% and 43.5%, respectively) (**Fig. 1b**). This improvement is likely attributable to Bst3’s enhanced enzymatic properties, including higher fidelity^17^, strong strand displacement activity, and optimal activity at elevated temperatures. We previously leveraged Bst3 in random priming step for single-cell joint-profiling of DNA methylation (5mC) and hydroxymethylation (5hmC)^18^, where it similarly improved alignment rates.

Lastly, we directly compared the optimized 2^nd^ SS protocol (Bst3 and N3G2N4B oligo) with the original method using Klenow DNA polymerase coupled with N9 oligo. The optimized method led to improved alignment rates (**Fig. 1d**), increased library complexity (**Fig. 1e**), and reduced background mutation rates (**Fig. 1f**).

Together, these results demonstrate that the combination of Bst3 DNA polymerase with a redesigned N3G2N4B primer significantly enhanced the performance of 2^nd^ SS reaction in scNT-seq analysis. We refer to this updated and optimized scNT-seq workflow as scNT-seq2 (**Fig. 2a**).

**Figure 2.**
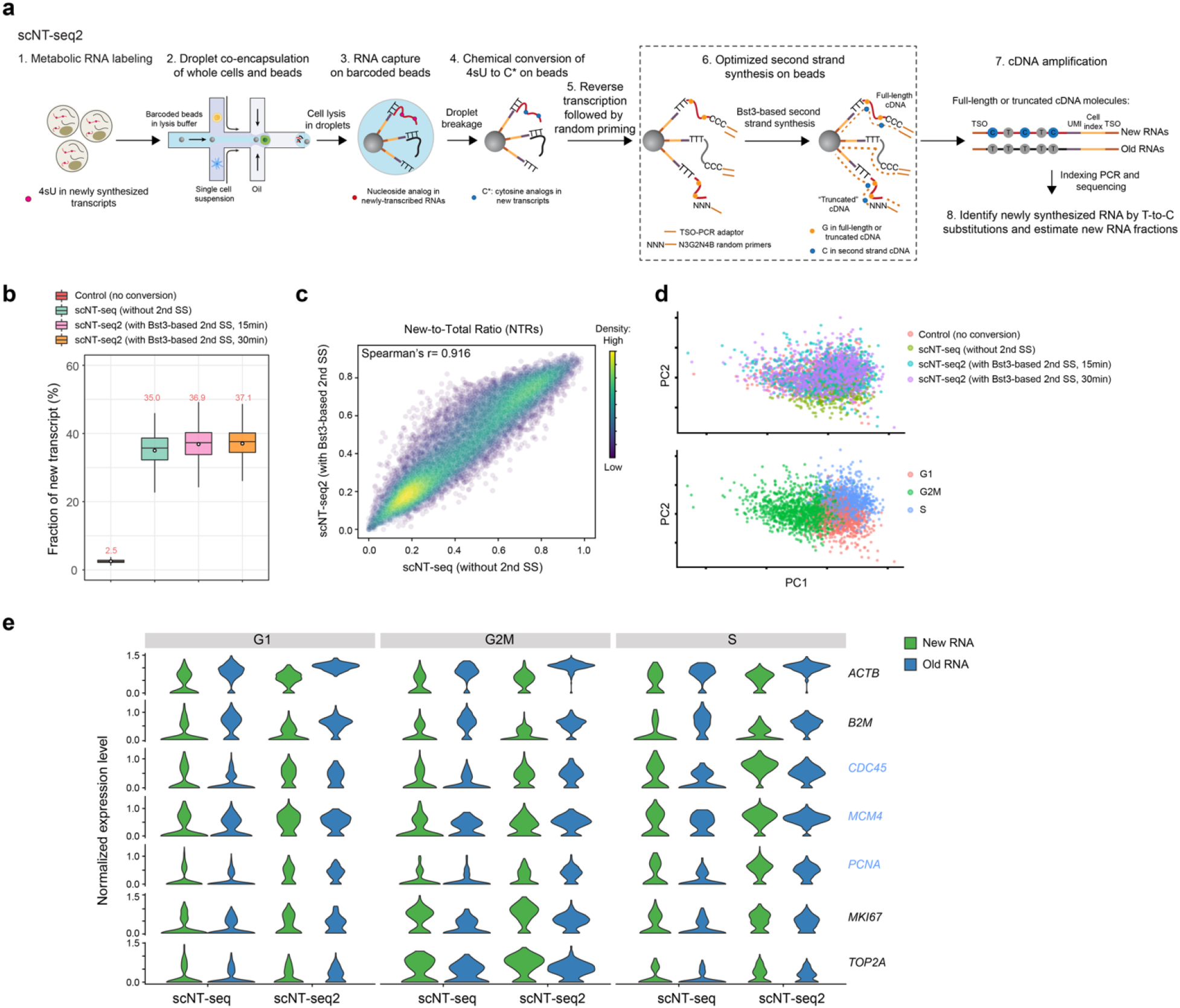
scNT-seq2 enhances the detection sensitivity of cell-cycle genes. **a**. Schematic depiction of the workflow of scNT-seq2. **b**. Box plot showing the proportion of metabolically labeled newly-synthesized transcripts per cell in K562 cells across different protocols with or without 2^nd^ SS. **c**. Scatterplots showing Spearman’s correlation for new and old RNA abundances and NTRs between original scNT-seq (without 2^nd^ SS) and scNT-seq2 (with Bst3-based 2^nd^ SS) methods. **d**. PCA plots showing K562 cells colored by experiments (top) or cell-cycle states (bottom). **e**. Violin plots showing the new and old RNA abundances of one house-keeping gene (*ACTB*) and 6 representative cell-cycle genes across three cell-cycle phases (G1/S/G2M). Three S-phase genes (*CDC45, MCM4*, and *PCNA*) are highlighted in blue.

### scNT-seq2 enhances the detection sensitivity of cell-cycle genes

To evaluate the performance of scNT-seq2, we benchmarked key metrics in metabolically labeled K562 cells across protocols with and without 2^nd^ SS step. We first compared the fraction of newly synthesized RNA and found that samples with 2^nd^ SS exhibited a similar proportion of new transcripts (37.0%) as those without 2^nd^ SS (35.0%), with comparable cell-to-cell variability (**Fig. 2b**). In addition, the expression levels of new and old RNAs, as well as New-to-Total Ratios (NTRs), were highly concordant between the two conditions (**Fig. 2c**), indicating that the addition of 2^nd^ SS does not perturb the quantification of RNA turnover rates.

To further validate the utility of scNT-seq2, we assessed its sensitivity in detecting cell-cycle-phase–specific gene expression in K562 cells. Principal component analysis (PCA) based on canonical cell-cycle genes clearly resolved major cell-cycle phases across datasets generated with different protocols—including Drop-seq (control), scNT-seq without 2^nd^ SS, and scNT-seq2 (Bst3-based 2^nd^ SS, 15 or 30 min) (**Fig. 2d**). Notably, scNT-seq2 improved the detection sensitivity for key cell-cycle markers such as *CDC45* and *MCM4* in S-phase (**Fig. 2e**), consistent with the enhanced library complexity observed in scNT-seq2 datasets.

## Discussion

In this study, we present scNT-seq2, an improved approach for time-resolved single-cell RNA sequencing. By systematically optimizing the second-strand cDNA synthesis (2^nd^ SS) step, we significantly improved read alignment rates, reduced background mutations, and enhanced library complexity, particularly for genes involved in dynamic processes such as the cell cycle.

Benchmarking experiments in K562 cells demonstrated that scNT-seq2 maintains accurate quantification of newly synthesized and pre-existing RNAs. Importantly, scNT-seq2 improved the detection of cell-state–specific transcripts, such as S-phase genes, reflecting its enhanced sensitivity and library complexity. These improvements expand the utility of time-resolved RNA sequencing for studying transcriptomic dynamics at single-cell resolution.

Beyond scNT-seq, the optimized 2^nd^ SS reaction developed here has broad applicability to other transcriptomic platforms. For example, spatial transcriptomics methods like Slide-seq and Slide-seqV2 have similarly benefited from second-strand synthesis to boost library complexity. The higher fidelity, strand displacement activity, and thermostability of Bst3 polymerase— combined with rational primer design—offer a generalizable strategy to recover low-abundance or truncated transcripts in both single-cell and spatial contexts. Thus, the improvements introduced in scNT-seq2 can be readily adapted to enhance the performance of diverse single-cell and spatial transcriptomics profiling technologies.

Together, our results establish scNT-seq2 as a robust and sensitive platform for time-resolved transcriptomics and highlight the potential of 2^nd^ SS optimization to advance next-generation single-cell and spatial omics.

## METHODS

### Cell culture and metabolic RNA labeling

Human K562 cells (ATCC, CCL-243) were cultured in RPMI media supplemented with 10% Fetal Bovine Serum (Sigma, F6178). For *in vitro* metabolic RNA labeling, K562 media was replaced with media supplemented with 4sU (100 μM). After 4 hours, the K562 cells were collected by spinning down, rinsed once with PBS, and fixed with methanol as described below.

### Cell fixation and rehydration for scNT-seq2

The cell fixation method was performed as previously described ^1,19^. Tissues were dissociated into single cell suspension as mentioned above. After resuspending with 0.4 mL of DPBS+ 0.01% BSA, the cells were split to two 1.5 mL LoBind tubes (Eppendorf) and added 0.8 mL methanol dropwise at a final concentration of 80% methanol in DPBS. After incubating the cell suspension on ice for 1 hour, store the fixed cells in LoBind tubes at -80°C freezer for up to one month. For rehydration, the cells were spun-down at 1000 g for 5 min at 4°C. Methanol-PBS solution was removed and cells were resuspended in 1 mL 0.01% BSA in DPBS supplemented with 0.5% RNase-inhibitor (Lucigen, 30281-2). After counting the cell number with Countess II system, the cell suspension was diluted to 100 cells/μL and immediately used for the scNT-seq2 experiment.

### Development and benchmark of second strand synthesis reaction using *in vitro* labeled human K562 cells

Human K562 cells were metabolically labeled with 4sU for 4 hours and subsequent fixed as detailed in the previous section. To further improve the performance of the second strand synthesis reaction, a series of random oligos were designed (**Supplementary Table 1**) and tested using previous work as references ^1,15^. In the random primer comparison experiment (**Supplementary Fig. 1b-c**), each oligo was added to a Klenow-based reaction mixture (200 μl of reaction mixture of Klenow buffer A consists of: 1x maxima RT buffer (ThermoFisher), 12% PEG-8000, 1 mM dNTPs (Clontech), 5 μM Template Switch Oligo-GAATG (TSO-GAATG: /5SpC3/AAGCAGTGGTATCAACGCAGAGTGAATG), 10 μM Template Switch Oligo-random primer, and 1.25 U/μl Klenow exo-(Enzymatics)) and incubated the beads for 60 min at 37°C with end-over-end rotation ^15^.

To improve the performance of scNT-seq2, we also tested different enzymatic reaction conditions. First, we tested the Klenow exo-enzyme (Enzymatics) in two different reaction buffers (Klenow buffer A: see above; buffer B: 200 μl of reaction mixture consists of: 1× Blue buffer (Enzymatics), 4% Ficoll PM-400, 1 mM dNTPs (Clontech), 10 μM Template Switch Oligo-random primer, and 1.25 U/μl Klenow exo-(Enzymatics)) (Qiu et al., 2020). For the Klenow-based reaction, the reaction was incubated for 60 min at 37°C with rotation. Second, we evaluated the *Bst* 3.0 DNA polymerase (denoted as Bst3; Bst3 reaction buffer: 1× Isothermal Amplification Buffer II (NEB), 6 mM MgSO4, 4% Ficoll PM-400, 1.4 mM dNTPs (Clontech), 10 μM Template Switch Oligo-random primer, and 0.4 U/μl *Bst* 3.0 DNA polymerase (NEB, M0374)), a DNA polymerase with strong strand displacement activity and higher fidelity than Klenow (**Supplementary Fig. 1d-f**). For the Bst3-based reaction, the reaction was incubated on ice for 2 min for primer annealing, followed by incubation at 60 °C for 15 min with rotation.

After systematic evaluation of the performance (i.e. library complexity, uniquely mapped reads percentage, and nucleotide substitution rates) with various random primers and DNA polymerase reactions, we identified the combination of TSO-N3G2N4B oligo with *Bst* 3.0 DNA polymerase as the most optimal conditions for the second strand synthesis reaction in scNT-seq2.

### scNT-seq2 library preparation and sequencing

The scNT-seq2 libraries were generated using the custom-built droplet microfluidics based scRNA-seq platform ^1^ with an extensively optimized second strand synthesis reaction (see above). Specifically, the single cell suspension was diluted to a concentration of 100 cells/μL in DPBS containing 0.01% BSA. Approximately 1.5 mL of diluted single cell suspension was loaded for each scNT-Seq2 run. The single-nucleus suspension was then co-encapsulated with barcoded beads (ChemGenes) using an Aquapel-coated PDMS microfluidic device (μFluidix) connected to syringe pumps (KD Scientific) via polyethylene tubing with an inner diameter of 0.38 mm (Scientific Commodities). Barcoded beads were resuspended in lysis buffer (200 mM Tris-HCl pH8.0, 20 mM EDTA, 6% Ficoll PM-400 (GE Healthcare/Fisher Scientific), 0.2% Sarkosyl (Sigma-Aldrich), and 50 mM DTT (freshly made on the day of run) at a concentration of 120 beads/μL. The flow rates for cells and beads were set to 3,200 μL/hour, while QX200 droplet generation oil (Bio-rad) was run at 12,500 μL/hour.

Droplet breakage with Perfluoro-1-octanol (Sigma-Aldrich). After droplet breakage, the beads were treated with TimeLapse reaction to convert 4sU to cytidine-analog ^3^. Briefly, 50,000-100,000 beads were washed once with 450 μL washing buffer (1 mM EDTA, 100 mM sodium acetate (pH 5.2)), then the beads were resuspended with a mixture of TFEA (600 mM), EDTA (1 mM) and sodium acetate (pH 5.2, 100 mM) in water. A solution of 192 mM NaIO_4_ was then added (final concentration: 10 mM) and incubated at 45°C for 1 hr with rotation. The beads were washed once with 1 mL TE, then incubated in 0.5 mL 1 X Reducing Buffer (10 mM DTT, 100 mM NaCl, 10 mM Tris pH 7.4, 1 mM EDTA) at 37°C for 30 min with rotation, followed by washing once with 0.3 mL 2X RT-buffer.

For reverse transcription (RT), up to 120,000 beads, 200 μL of RT mix (1x Maxima RT buffer (ThermoFisher), 4% Ficoll PM-400, 1 mM dNTPs (Clontech), 1 U/μL RNase inhibitor, 2.5 μM Template Switch Oligo (TSO: AAGCAGTGGTATCAACGCAGAGTGAATrGrGrG) ^20^, and 10 U/μL Maxima H Minus Reverse Transcriptase (ThermoFisher)) were added. The RT reaction was incubated at room temperature for 30 minutes, followed by incubation at 42°C for 120 minutes. After exonuclease I treatment, an optimized second strand synthesis reaction was applied to improve library complexity and alignment rate (see above). Specifically, pooled beads were washed once with TE-SDS buffer and twice with TE-TW buffer to remove the exonuclease I reaction mixture. The beads were resuspended in 500 μl of 0.1 M NaOH and incubated for 5 min at room temperature with rotation, and 500 μl of 0.2 M Tris-HCl (pH 7.5) was then added to neutralize the solution. The beads were washed once with TE-TW buffer and once with 10 mM Tris-HCl (pH 8.0). To synthesize second strand cDNA, the beads were resuspended in 200 μl of *Bst*3-based reaction mixture (1× Isothermal Amplification Buffer II (NEB), 6 mM MgSO4, 4% Ficoll PM-400, 1.4 mM dNTPs (Clontech), 10 μM Template Switch Oligo-random primer (TSO-N3G2N4B:AAGCAGTGGTATCAACGCAGAGTGA (N1:25252525)(N1)(N1)GG(N1)(N1)(N1) (N1)(N2: 00333433); N1 represents a mixture of A, C, G and T at a 25:25:25:25 ratio, N2 represents a mixture of A, C, G and T at a 0:33:34:33 ratio), and 0.4 U/μl *Bst* 3.0 DNA polymerase (NEB, M0374)). The reaction was incubated on ice for 2 min for primer annealing, followed by incubation at 60 °C for 15 min with rotation. The reaction was stopped by washing the beads once with TE-SDS buffer and twice with TE-TW buffer.

The downstream steps (cDNA amplification, Tn5-based tagmentation, and indexing PCR) were performed as previously described for scNT-seq ^1^. Briefly, to determine an optimal number of PCR cycles for amplification of cDNA, an aliquot of 6,000 beads was amplified by PCR in a volume of 50 μL (25 μL of 2x KAPA HiFi hotstart readymix (KAPA biosystems), 0.4 μL of 100 μM TSO-PCR primer (AAGCAGTGGTATCAACGCAGAGT, 24.6 μL of nuclease-free water) with the following thermal cycling parameter (95°C for 3 min; 4 cycles of 98°C for 20 sec, 65°C for 45 sec, 72°C for 3 min; 9 cycles of 98°C for 20 sec, 67°C for 45 sec, 72°C for 3 min; 72°C for 5 min, hold at 4°C). After two rounds of purification with 0.6x SPRISelect beads (Beckman Coulter), amplified cDNA was eluted with 10 μL of water. 10% of amplified cDNA was used to perform real-time PCR analysis (1 μL of purified cDNA, 0.2 μL of 25 μM TSO-PCR primer, 5 μL of 2x KAPA FAST qPCR readymix, and 3.8 μL of water) to determine the additional number of PCR cycles needed for optimal cDNA amplification (Applied Biosystems QuantStudio 7 Flex). We then prepared PCR reactions per total number of barcoded beads collected for each scNT-seq2 run, with ∼6,000 beads per PCR tube, and ran the PCR program to enrich the cDNA for 4 plus 10-12 cycles. We then tagmented cDNA using the Nextera XT DNA sample preparation kit (Illumina, cat# FC-131-1096), starting with 550 pg of cDNA pooled in equal amounts, from all PCR reactions for a given run. Following cDNA tagmentation, we further amplified the library with 12 enrichment cycles using the Illumina Nextera XT i7 primers along with the P5-TSO hybrid primer ^20^. After quality control analysis using a Bioanalyzer (Agilent), libraries were sequenced on an Illumina NextSeq 500 instrument using the 75-cycle High Output v2.5 Kit (Illumina). We loaded the library at 2.0 pM and provided Custom Read1 Primer (GCCTGTCCGCGGAAGCAGTGGTATCAACGCAGAGTAC) at 0.3 μM in position 7 of the reagent cartridge. The sequencing configuration was 20 bp (Read1), 8 bp (Index1), and 60 bp (Read2).

### scNT-seq2 data preprocessing

The preprocessing of scNT-seq2 datasets, encompassing read alignment, quality filtering, and count matrix generation, was performed utilizing the Dynast pipeline (version 1.0.1; available at https://github.com/aristoteleo/dynast-release), which is an inclusive and efficient toolkit for preprocessing metabolic labeling-based scRNA-seq experiments, with modifications specific to the droplet microfluidics based scNT-seq2. The Dynast pipeline is structured as a multi-step workflow consisting of ‘dynast align’, ‘dynast consensus’, ‘dynast count’, and ‘dynast estimate’. First, raw sequencing reads were mapped to the human genome GRCh38 (genecode v41 annotation) using ‘dynast align’ with the paired-end mode (i.e., “--strand forward -x dropseq”). Cells were filtered with the parameter “--soloCellFilter TopCells” from ‘STARsolo’ to only retain high quality cells for all downstream analyses. Second, ‘dynast consensus’ involved constructing a consensus sequence for each transcript pooling reads with the same UMI index and taking the most frequent variant at each position as previously reported ^1^. ‘dynast count’ was employed to quantify the levels of unspliced, spliced, newly-synthesized, and total RNA for each single cell. In addition, sites with background T-to-C substitutions (present in the control sample without TimeLapse-based chemical conversion treatment) were determined and excluded for T-to-C substitution identification.

### Cell-type clustering and marker gene identification for scNT-seq2

All downstream data preprocessing, clustering, and differential expression analysis steps were performed using Scanpy (v1.9.1). The RNA count matrices were then imported into Scanpy as AnnData objects. The analysis pipeline for scNT-seq2 datasets consists of following main steps:

#### Quality control

Cells with fewer than 500 or more than 10,000 genes, or more than 10% mitochondrial genes were filtered out. For all scNT-seq2 datasets, genes expressed in fewer than 3 cells were also removed.

#### Clustering and Visualization

After preprocessing, the data was clustered and visualized using the following procedure. Cell cycle scores were calculated with Seurat (v4.1.1) using *CellCycleScoring* function. Principal component analysis (PCA) on cell cycle genes was performed to separate K562 cells by cell cycle phase (**Fig. 2d**).

## Materials Availability

The experimental protocol generated in this study is available via protocol.io (https://www.protocols.io/view/scnt-seq2-single-cell-metabolically-labelled-new-r-j8nlk8811l5r/v1).

## Data availability

All sequencing data associated with this study will be available on the NCBI Gene Expression Omnibus (GEO) database upon publication.

## Code availability

The analysis source code underlying the final version of the paper will be available on GitHub repository (https://github.com/wulabupenn/) upon publication.

## ACKNOWLEDGEMENTS

We are grateful to all members of the Wu lab for helpful discussion. This work is in part supported by a National Human Genome Research Institute (NHGRI) grant U01HG012047 (to H.W.), a National Institute of Neurological Disorders and Stroke (NINDS) grant U19NS135528 (to H.W.), and a National Eye Institute (NEI) grant U01EY034681 (to H.W.).

## AUTHOR CONTRIBUTIONS

Q.Q., and H.W. conceived the study. Q.Q. conducted all experiments and most of the data analysis with help from H.Z. and W.G.. Q.Q., W.G., and H.W. interpreted the data and wrote the manuscript with contributions from all authors.

## Supplementary Figure and Tables

**Supplementary Figure 1.**
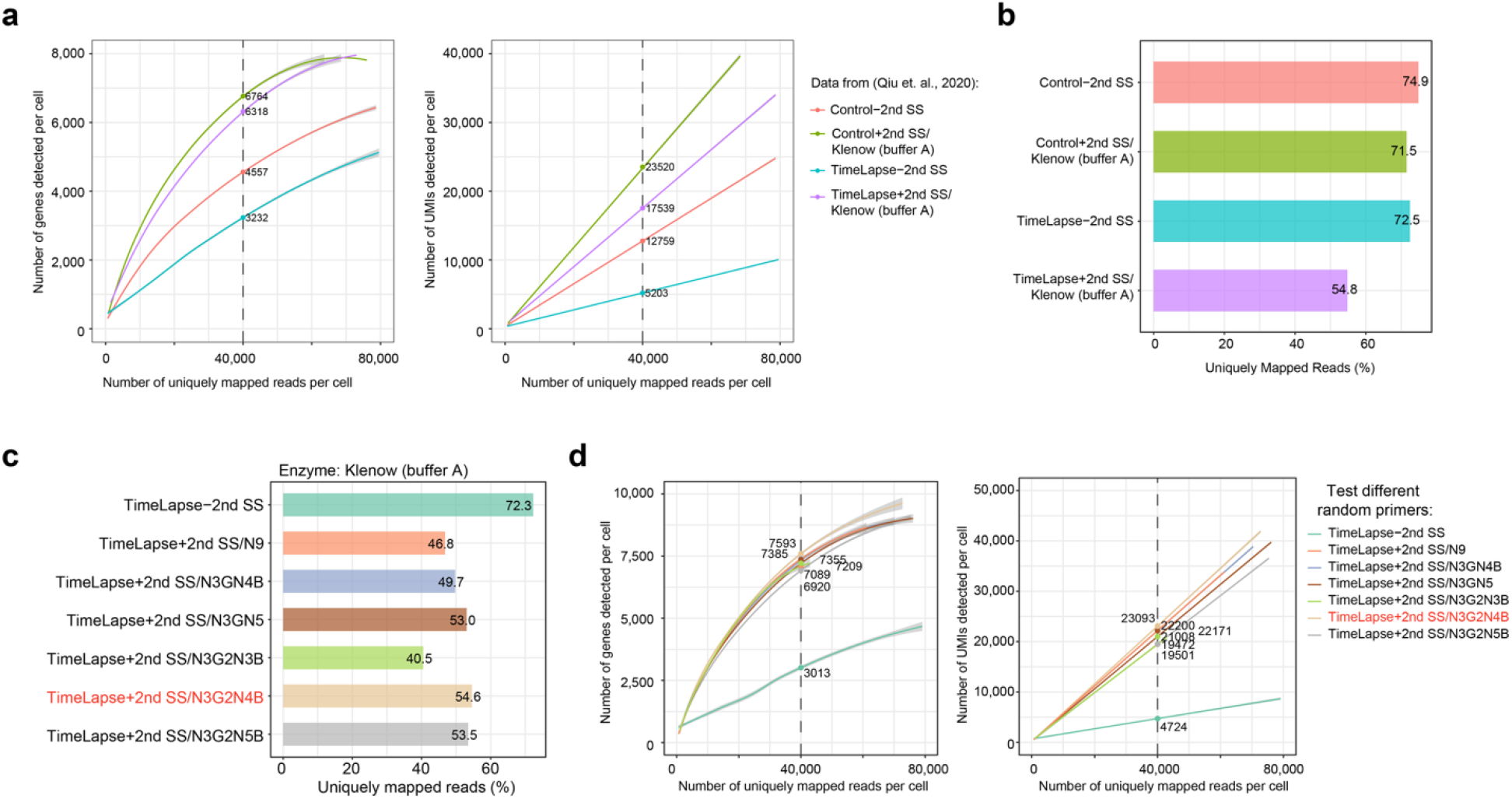
Benchmarking the Klenow-based second-strand cDNA synthesis reactions. **a**. Fitted line plots illustrating library complexity by comparing genes (left) or UMIs/transcripts (right) detected per cell as a function of aligned reads per cell across different conditions using the data from Qiu et. al., 2020 ^1^. All experiments were performed using the same batch of *in vitro* 4sU-labeled K562 cells (100 μM, 4 hours). Estimated numbers of genes or UMIs detected per cell at matching sequencing depth (40,000 reads per cell) for different experiments are shown. The shaded regions depict 95% confidence intervals. **b**. Bar graph showing the fraction of uniquely mapped reads various different experiments in (**a**). **c**. Fitted line plots illustrating library complexity by comparing genes (left) or UMIs/transcripts (right) detected per cell as a function of aligned reads per cell across experiments with various random primers. This set of experiments used a previously reported Klenow-based reaction mixture ^15^, with different random primer sequences (provided in **Supplementary Table 2**). All experiments were performed using the same batch of *in vitro* 4sU-labeled K562 cells (100 μM, 4 hours). Estimated numbers of genes or UMIs detected per cell at matching sequencing depth (40,000 reads per cell) for different experiments are shown. The shaded regions depict 95% confidence intervals. **d**. Bar graph showing the fraction of uniquely mapped reads for the experiments in (**d**).

**Supplementary Table 1.**
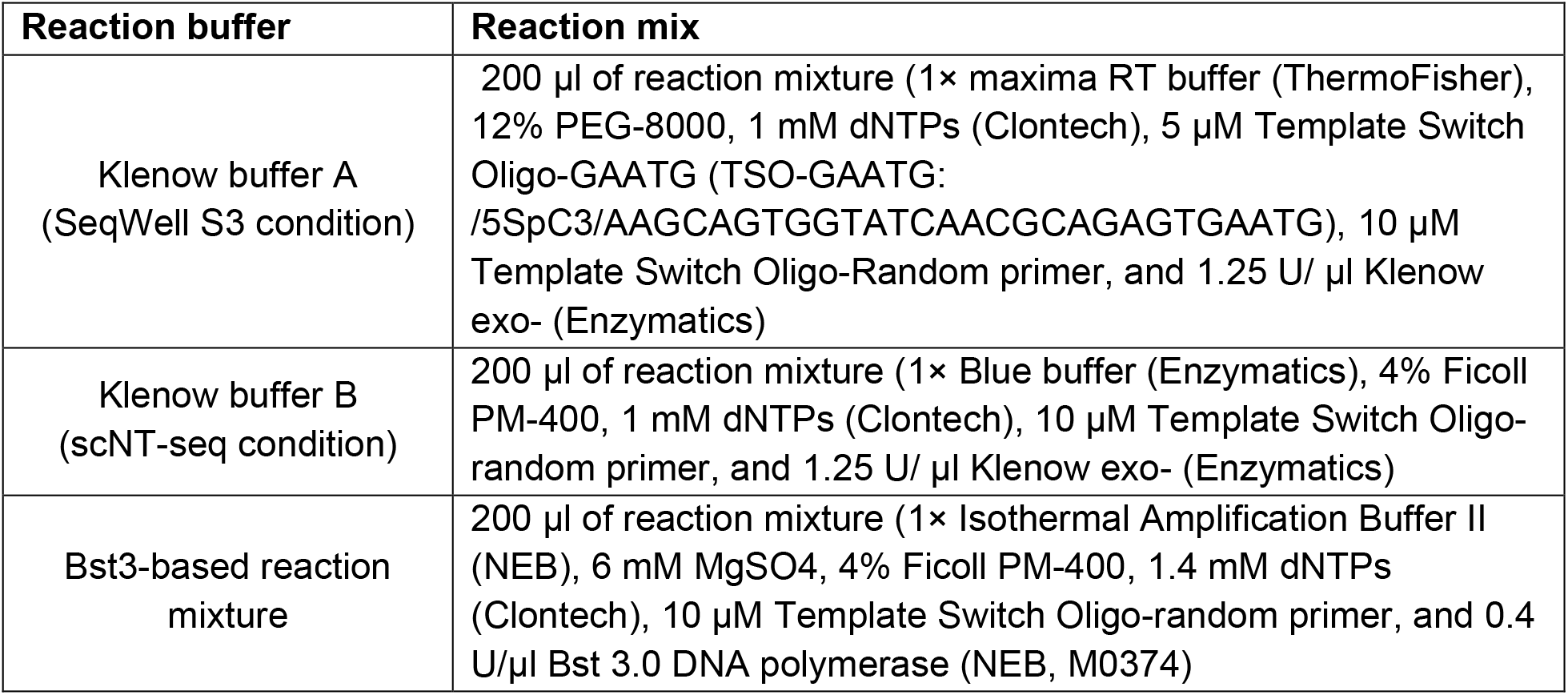

**Supplementary Table 2.**
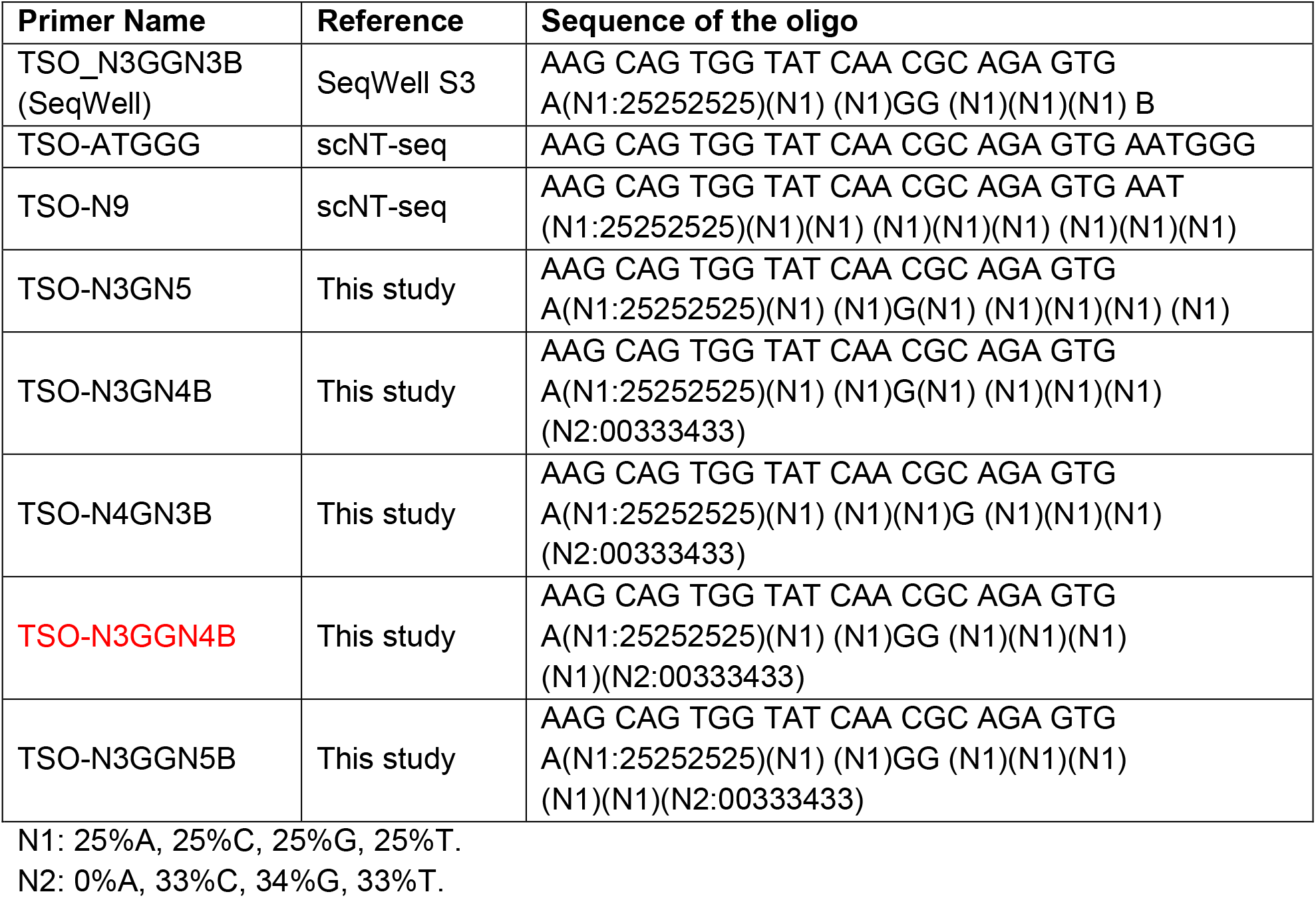

